# Multi-omic Profiling Reveals Early Immunological Indicators for Identifying COVID-19 Progressors

**DOI:** 10.1101/2023.05.25.542297

**Authors:** Katherine A. Drake, Dimitri Talantov, Gary J. Tong, Jack T. Lin, Simon Verheijden, Samuel Katz, Jacqueline M. Leung, Benjamin Yuen, Vinod Krishna, Michelle J. Wu, Alex Sutherland, Sarah A. Short, Pouya Kheradpour, Maxwell Mumbach, Kate Franz, Vladimir Trifonov, Molly V. Lucas, James Merson, Charles C. Kim, PRESCO Study Group

## Abstract

The pandemic caused by the severe acute respiratory syndrome coronavirus 2 (SARS-CoV-2) has led to a rapid response by the scientific community to further understand and combat its associated pathologic etiology. A focal point has been on the immune responses mounted during the acute and post-acute phases of infection, but the immediate post-diagnosis phase remains relatively understudied. We sought to better understand the immediate post-diagnosis phase by collecting blood from study participants soon after a positive test and identifying molecular associations with longitudinal disease outcomes. Multi-omic analyses identified differences in immune cell composition, cytokine levels, and cell subset-specific transcriptomic and epigenomic signatures between individuals on a more serious disease trajectory (Progressors) as compared to those on a milder course (Non-progressors). Higher levels of multiple cytokines were observed in Progressors, with IL-6 showing the largest difference. Blood monocyte cell subsets were also skewed, showing a comparative decrease in non-classical CD14^−^CD16^+^ and intermediate CD14^+^CD16^+^ monocytes. Additionally, in the lymphocyte compartment, CD8^+^ T effector memory cells displayed a gene expression signature consistent with stronger T cell activation in Progressors. Importantly, the identification of these cellular and molecular immune changes occurred at the early stages of COVID-19 disease. These observations could serve as the basis for the development of prognostic biomarkers of disease risk and interventional strategies to improve the management of severe COVID-19.

**One Sentence Summary:** Immunological changes associated with COVID-19 progression can be detected during the early stages of infection.

## INTRODUCTION

Since the beginnings of the COVID-19 pandemic, the immune response has been intensely studied in patients from the standpoint of both the host antiviral response and unchecked inflammatory pathology *(1, 2)*. While this has resulted in the rapid development of testing, prophylactic, and interventional methods for managing the spread and severity of SARS-CoV-2 infection *(3)*, the health burden of resultant COVID-19 remains high. Global excess deaths from the period of January 2020 to December 2021 were estimated to be 14.83 million, with the top 20 countries accounting for over 80% of deaths *(4)*. Even at the beginning of 2023, 7-day rolling averages of excess deaths due to COVID-19 were still 350+ in the United States and 1,850+ worldwide *(5)*. If this pace continues without abatement, mortality from COVID-19 in the United States will be on track to continue to exceed that of both influenza and respiratory syncytial virus, the two most common causes of respiratory infection-associated deaths before the pandemic, combined *(6)*. However, it has also been recognized that the majority of individuals infected with SARS-CoV-2 develop a relatively mild, self-resolving COVID-19 disease that does not require medical intervention, and it is only in a minority of patients that a severe disease course occurs requiring hospitalization and costly interventions *(7–9)*. This heterogeneity, along with continuing mortality, highlights the need for a deeper understanding of COVID-19 pathogenesis.

Since the mechanisms behind this difference in disease progression are not yet well understood, multi-omic approaches that provide a broad and high resolution overview of immune function can help form hypotheses about the cellular interplay and affected pathways that preceded and shaped the current immune state. Such multi-omic studies have previously been carried out in COVID-19 patients, with much of the focus on the mechanisms and signatures of severe disease at later phases in infection *(10–17)*, and also with confounding influence of vaccines and oral antivirals. Indeed, many of these severe disease signatures such as immune cell activation and exhaustion *(12, 18, 19)*, myeloid cell dysfunction *(12, 15, 17, 20)*, elevated inflammatory cytokines and impaired type-I interferon production *(11–13, 21, 22)* have since been targets for COVID-19 clinical trials *(23–28)*. Some success has been achieved through targeting the immune system with immune-modulating therapeutics including infliximab (anti-TNF antibody) *(23)*, tocilizumab (anti-IL-6R antibody) *(23, 24)*, anakinra (IL-1R antagonist) *(25)*, and abatacept (co-stim inhibitor CTLA4-Ig) *(26)*, which improved survival in severe COVID-19 patients.

Our study sought to characterize the immune responses to COVID-19 from biospecimens collected during the early phase of SARS-CoV-2 infection using multi-omic assays. We compared early immune response signatures in participants classified as Progressors (those requiring hospitalization or outpatient treatment for COVID-19 or for the worsening of a pre-existing condition triggered by SARS-CoV-2 infection within 28 days) and Non-progressors (those who did not require such clinical interventions for their COVID-19 disease within 28 days) to find mechanisms that distinguish those on higher risk trajectories from milder disease. Findings from our analyses corroborate aspects of the altered immune response described in later infection, but demonstrate that many molecular and cellular changes occur during early infection.

## RESULTS

### Study population and molecular data generation

We conducted the COVID Progression Retrospective (CPR) study, which was a retrospective cross-sectional study of existing datasets and biospecimens from two COVID-19 studies (COVID-19 Immune Response Study (Cove) and Predictors of Severe COVID-19 Outcomes (PRESCO; clinicaltrials.gov NCT04388813)), with the aim of capturing the earliest immunological signature of disease progression upon confirmed SARS-CoV-2 infection. A total of 198 participant datasets were eligible for the CPR study. Of these, 14 participant datasets were excluded from analysis due to insufficient information in determining a progression outcome. Additionally, 22 participant datasets were excluded from analysis because they: 1) did not meet requirements for study completion, 2) had evidence of some immunomodulating treatments prior to biospecimen collection, or 3) did not report any symptoms prior to or throughout the study. Lastly, 4 additional participant datasets were excluded from analysis due to lack of biospecimen availability and/or provenance. In total, 162 participant datasets were eligible for descriptive analyses of demographic and clinical variables, and 158 participant datasets were included for the molecular analyses in the CPR study.

Demographic and clinical characteristics of the analysis population were summarized (**Table 1**). Additional participant characteristics included body mass index (BMI), smoking frequency and years of use, and comorbidities classified by affected physiologic systems (**table S1**). Upon analyzing these demographics and clinical characteristics, we found advanced age, female sex, Black race, high BMI, and presence of comorbidities including cardiovascular, metabolic, renal, and respiratory conditions to be significant predictors of progression (**table S2**). With the exception of female sex, these observations are consistent with previously identified and reported associations with severe COVID-19 *(29–33)*.

**Table 1.** Demographics of study participants.

| **Characteristics** | **Total**<br>**(N = 162)** | **Progressor**<br>**(N = 24)** | **Non-progressor**<br>**(N = 138)** |
| --- | --- | --- | --- |
| Age, mean (SD) | 41.4 (14.8) | 53 (18) | 39 (13) |
| Age category, N (%) |  |  |  |
| 18-29 | 41 (25.3%) | 2 (8.3%) | 39 (28.3%) |
| 20-49 | 75 (46.3%) | 10 (41.7%) | 65 (47.1%) |
| 50-64 | 35 (21.6%) | 6 (25%) | 29 (21%) |
| 65+ | 11 (6.8%) | 6 (25%) | 5 (3.6%) |
| Female, N (%) | 98 (60.5%) | 20 (83.3%) | 78 (56.5%) |
| Race*, N (%) |  |  |  |
| White | 87 (62.1%) | 8 (36.4%) | 79 (66.9%) |
| Black | 45 (32.1%) | 13 (59.1%) | 32 (27.1%) |
| Asian | 7 (5%) | 0 (0%) | 7 (5.9%) |
| American Indian / Alaskan Native | 2 (1.4%) | 1 (4.5%) | 1 (0.8%) |
| Native Hawaiian / Pacific Islander | 1 (0.7%) | 0 (0%) | 1 (0.8%) |
| Race unknown / not reported, N (%) | 22 (13.6%) | 2 (8.3%) | 20 (14.5%) |
| Hispanic*, N (%) | 39 (24.4%) | 5 (20.8%) | 34 (25%) |
| Ethnicity unknown / not reported, N (%) | 2 (1.2%) | 0 (0%) | 2 (1.4%) |

*\* Percentages calculated out of participants with non-missing data*

Participant blood samples and other biospecimens had an overall mean time to collection of 3.2 days post-positive test. Peripheral blood from 158 participants was processed through a multi-omic immunogenomic profiling workflow, consisting of immune cell subset sorting by fluorescence-activated cell sorting (FACS), followed by RNA sequencing (RNA-seq), assay for transposase-accessible chromatin using sequencing (ATAC-seq), targeted protein estimation by sequencing (TaPE-seq), and whole genome sequencing (WGS). The 24 immune cell subsets resolved by fluorescent surface marker staining and sorting represent canonically defined subsets with respect to cell surface markers and known characteristics of lineage and functional differentiation (**table S3**). An additional sorted bulk peripheral blood mononuclear cell (PBMC) subset was used as a quality control metric and was not further analyzed. Over 8 million molecular features generated for each participant PBMC sample were analyzed to identify differences between Progressors and Non-progressors, along with measures of viral load from mid-turbinate swabs and plasma cytokine levels (**table S4**). There were no significant differences in the viral load between Progressors and non-Progressors early in the disease course, as measured by detection of the viral genes *N* and *ORF1a* by ddPCR or in the IgG antibody reactivity towards SARS-CoV-2 S and N protein or other related human coronaviruses (**fig. S1**).

### Progression is associated with early changes in monocyte and T cell subsets

Progressor and Non-progressor cell subset frequencies (measured as a percentage of PBMCs) across the 24 subsets were assessed, and no significant differences were observed after adjusting for multiple testing (Fig. 1A), suggesting that early timepoints of progression are not correlated with cell frequency shifts when viewed across the global PBMC population. To assess whether there are differences in coordinated immune responses early in infection, a pairwise correlation analysis of cell subset frequencies was conducted (**fig. S2**). Progressors and Non-progressors showed some clustering of myeloid and B cell subsets, while an effector T cell cluster was seen only in Progressors.

**Fig. 1.**
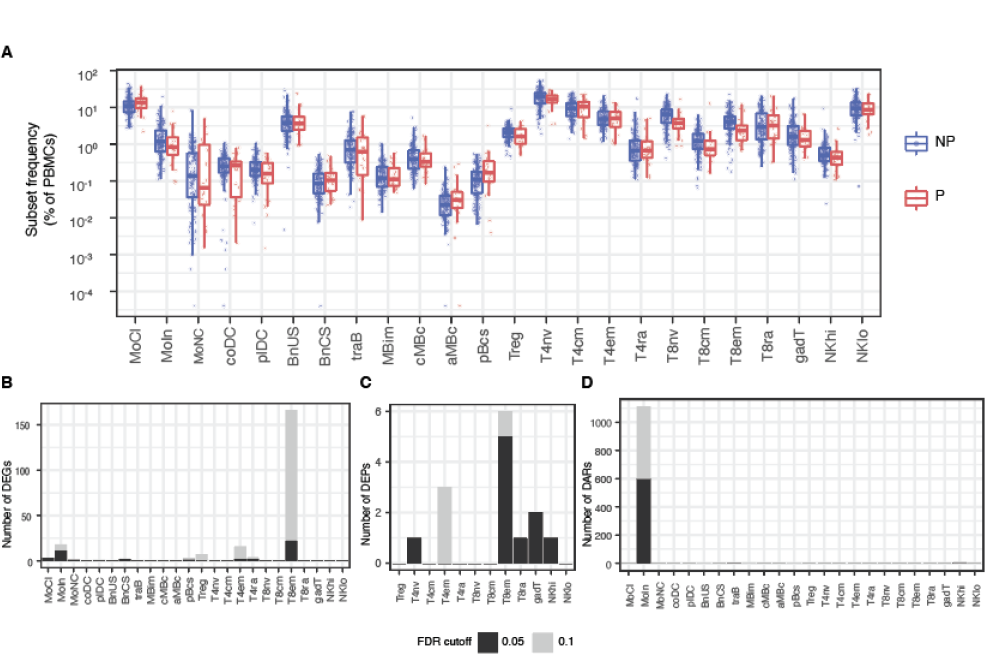
Immune Profiler cell frequency and subset-resolved multi-omic data overview. (**A**) Box plots of cell subset frequencies grouped by Non-progressor (NP) versus Progressor (P) pairwise outcome groups. (**B**) Identified number of significant differentially expressed genes (DEGs) between Non-progressors and Progressors by cell subset from RNA-seq data. (**C**) Identified number of differentially expressed proteins (DEPs) between Non-progressors and Progressors by cell subset from TaPE-seq data. (**D**) Identified number of differentially accessible regions (DARs) between Non-progressors and Progressors by cell subset from ATAC-seq data; For bar graphs (B to D), FDR cutoffs for each data type were set and graphed as FDR ≤ 0.05 (black bar) or FDR ≤ 0.1 (gray bar); Abbreviations for cell subsets are defined in table S3.

Differential expression analysis identified transcripts in specific cell subsets up- or down-regulated in Progressors (Fig. 1B). The highest number of differentially expressed genes (DEGs) was seen in the effector memory CD8^+^ T cell subset, followed by the intermediate monocyte subset. Pathway analysis using Gene Set Enrichment Analysis (GSEA) and Gene Set Variation Analysis (GSVA) showed the highest number of significant pathways in the effector memory CD8^+^ T cell subset (**fig. S3, A and B**). Assessment of cell surface protein expression on the T cell and NK cell subsets revealed the highest number of differentially expressed proteins (DEPs) in the effector memory CD8^+^ T cell subset (Fig. 1C). The majority of differentially accessible regions (DARs) identified with ATAC-seq were observed in the intermediate monocyte subset (Fig. 1D, **fig. S3C**).

Multi-omic approaches that integrate “regulatory layer” evidence from functional genomic or genetic assays can build confidence in DEGs identified from differential expression analysis by linking them to genetic factors. To this end, we employed an approach to identify expression quantitative trait loci (eQTLs) that correspond to the DEGs identified from differential analysis. We generated cell subset-resolved cis-eQTLs and observed that DEGs (at false discovery rate (FDR) ≤ 0.1) in effector memory CD8^+^ T cells and intermediate monocytes were also subset-resolved cis-eQTLs (**table S5**).

Finally, multi-omic machine learning was used to identify cell subsets with the most discriminative features for classifying progression outcome. Block sparse partial least squares-discriminant analysis was performed for each cell subset utilizing gene expression, chromatin accessibility, and progression outcomes *(34)*. This analysis showed intermediate monocytes, naive CD4^+^ T cells, and effector memory CD8^+^ T cells as the top three cell subsets whose RNA and ATAC features best classified progression outcome (**fig. S3D**).

In summary, this high-resolution multi-omic approach revealed significant immune signatures already differentiating Progressors from Non-progressors during early SARS-CoV-2 infection, before clinical intervention. Understanding what cell subsets had the greatest molecular divergence between the two outcome groups helped guide further cell subset-specific analysis, with a focus on monocytes and effector memory CD8^+^ T cells.

### Early inflammatory cytokine and chemokine responses in Progressors

To determine the association between plasma cytokines and COVID-19 disease progression, we analyzed 44 total cytokines, chemokines, and growth factors in plasma during this early phase of SARS-CoV-2 infection. At an FDR ≤ 0.05, seven cytokines (IL-6, IFNγ, IL-7, PDGF-AA, TNF, MIG, and IL-15) were significantly higher in Progressors (Fig. 2A). At an FDR ≤ 0.1, 24 total cytokines were higher in Progressors (**fig. S4**), including key inflammatory cytokines such as IL-6, IP-10, IFNγ, TNF, and IL-18, as well as chemokines such as MCP-3 (CCL7), MIG (CXCL9), IP-10 (CXCL10), and Fractalkine (CX3CL1) (**fig. S4**).

**Fig. 2.**
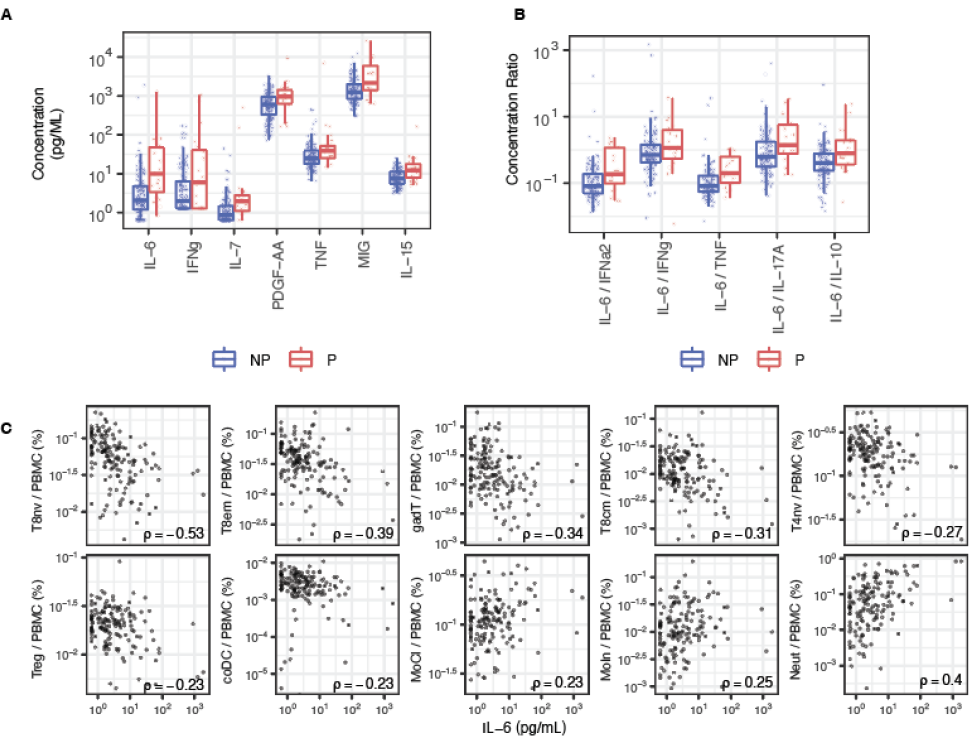
Elevated cytokine and chemokine levels in Progressors. (**A**) Concentration in pg/ml of cytokines and chemokines elevated in plasma of Non-progressor (NP) versus Progressor (P) at a FDR ≤ 0.05 as measured by Luminex assay. (**B**) Cytokine concentration ratios for NP vs P. (**C**) Spearman’s correlations between immune cell subset frequencies (as a percentage of PBMCs) and plasma IL-6 concentrations with a ρ ≥ 0.2 or ρ ≤ −0.2. Abbreviations for cell subsets are defined in table S3.

IL-6 was the cytokine with the largest difference in magnitude between Progressors and Non-progressors (Fig. 2A **and fig. S4**). As cytokines are often produced in a cascade, we evaluated various cytokine ratios to help discern the type of immune response and immune regulation occurring in COVID-19 progression. We computed ratios of IL-6 and common type I interferons (e.g., IFNα2) and Th1 (e.g., IFNγ and TNF), Th17 (e.g., IL-17A) and anti-inflammatory (e.g., IL-10) cytokines within individuals. For each cytokine ratio, the IL-6 response relative to the other common cytokines trended higher in Progressors compared to Non-progressors (Fig. 2B). We also examined cytokine gene expression levels in myeloid, B, and T cell subsets by RNA-seq to better disentangle the various contributions by cells to cytokine production (**fig. S5**). Focusing on the key inflammatory cytokine genes *IL6*, *IFNG*, and *TNF*, we found that expression of *IL6* trended higher in intermediate monocytes in Progressors, whereas expression of *IFNG* and *TNF* in Progressors trended higher in the CD4^+^ and CD8^+^ T cell subsets (**fig. S5**).

It was recently reported that plasma concentrations of IL-6, and to a lesser extent IL-10, correlated with increases in CD14^+^ monocytes with phenotypic similarity to myeloid-derived suppressor cells, which are capable of suppressing T cell proliferation *(35)*. We found that plasma IL-6 concentrations negatively correlated with the frequencies of a number of circulating T cell subsets, including subsets of conventional CD4^+^ and CD8^+^ T cells, regulatory T cells, and gamma delta T cells, and positively correlated with the frequencies of classical and intermediate monocytes and neutrophils (Fig. 2C).

### Cellular and molecular perturbations in the monocyte compartment of Progressors

We investigated relative frequencies measured by cytometry in the total monocyte population and found that the ratios of intermediate monocytes, non-classical monocytes, and CD16^+^ (intermediate + non-classical) monocytes over the total monocytes were all significantly lower in Progressors (p ≤ 0.01, p ≤ 0.05, p ≤ 0.001; respectively; Fig. 3A). By contrast, the ratio of classical monocytes over total monocytes was not significantly different between the two groups. Measuring chromatin accessibility of each subset further revealed differences in monocyte populations between the two outcome groups. Principal component analysis of accessible chromatin regions in the three monocyte subsets recapitulated the known developmental trajectory from classical, to intermediate, to non-classical, and showed a trend where Progressor intermediate monocytes grouped more closely with classical monocytes (Fig. 3B). The Progressor intermediate monocyte centroid was significantly closer to the classical monocyte centroid and further from the non-classical monocyte centroid in principal component space (Fig. 3C).

**Fig. 3.**
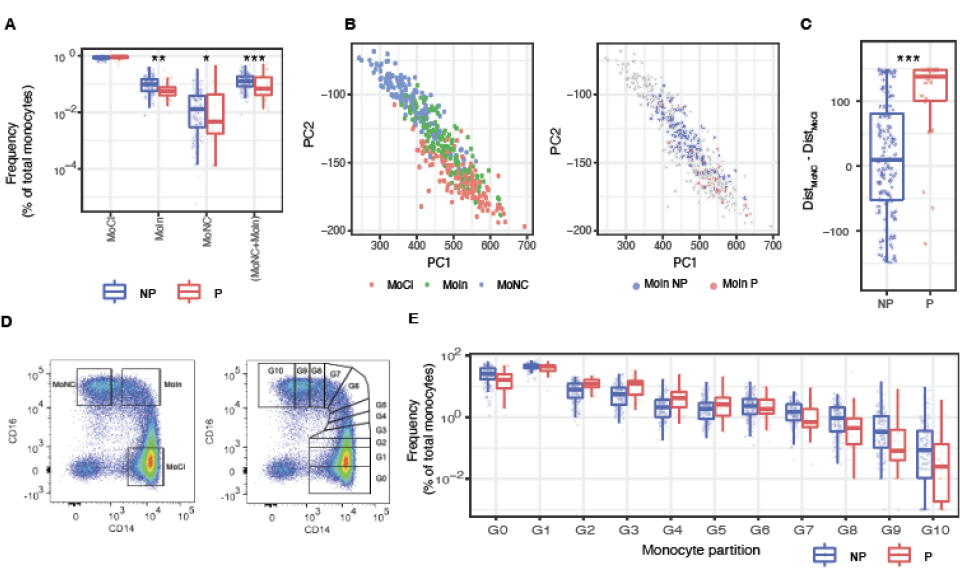
Perturbations in Progressor monocyte cell subset frequencies. (**A**) Ratio of monocyte subsets to total frequency of monocytes in Non-progressors (NP) versus Progressors (P). Significance assessed by p-values for each individual test. (**B**) Principal Component Analysis of all participant monocyte subsets’ ATAC-seq data, showing a spectrum ranging from classical monocytes (MoCl; red), to intermediate monocytes (MoIn; green), to non-classical monocytes (MoNC; blue) (left panel). The intermediate monocytes of NP (blue) or P (red) are colored (right panel). (**C**) Box plot of NP and P’s intermediate monocytes on the ATAC-Seq epigenetic spectrum. A Wilcoxon signed-rank test was performed to measure significant differences in Progressor versus Non-progressor intermediate monocytes based on their distances between the MoCl centroid and the MoNC centroid. (**D**) Typical biaxial CD14/CD16 plot showing defined gates for monocytes that delineate classical, intermediate, and non-classical monocyte subsets (left panel). Finer partitions along the biaxial CD14/CD16 plot, from G0-G10, allow for comparative measurements of frequency across the classical, intermediate, non-classical monocyte transitional path (right panel). (**E**) Cell frequencies of the G0-G10 gating partitions to total monocytes in NP versus P. Gating partitions are typically ascribed to classical (G0-G1), intermediate (G6-G7), and non-classical (G9-G10) monocyte CD14/CD16 gates; *p≤0.05; **p≤0.01; ***p≤0.001

To further investigate the observed differences in monocyte development and identity, we adapted a finer gating method for distinguishing monocyte subsets by flow cytometry using the cell surface markers CD14 and CD16. As previously described *(36)*, we partitioned monocytes into 10 partitions across CD14 and CD16 positions along the developmental path from CD14^+^CD16^−^ classical monocytes through CD14^+^CD16^+^ intermediate monocytes, to CD14^−^CD16^+^ non-classical monocytes (Fig. 3D). The frequencies of each gated partition showed a trend of increasing frequencies for Progressors as the gating moved from the classical to intermediate monocyte partitions (G2 to G5, Fig. 3E). This trend reversed upon reaching the CD14^+^CD16^+^ intermediate monocyte region with Progressors showing decreasing cell frequencies compared to Non-progressors in the partitions gated between intermediate and non-classical monocytes (G6 to G10, Fig. 3E). These findings are consistent with the model that circulating monocytes egress to sites of pathology, preventing the peripheral development of more mature intermediate and non-classical monocyte subsets.

The DEGs and DARs in monocytes further revealed differences between Progressors and Non-progressors. Monocytes of Progressors showed increased expression in genes and pathways that have been previously associated with later stage severe COVID-19 (Fig. 4A **and fig. S6, A and B**). At FDR ≤ 0.1, *CD163*, *CLU*, *LYN*, *MCEMP1*, *RAB13*, *RNASE1*, and *THBD* were found to be differentially expressed between the two groups and were also proximal to DARs (Fig. 4B). Previous reports have identified *CD163* expression in monocytes and soluble CD163 in the serum as being correlated with disease *(37–39)*, and *MCEMP1* and *THBD* have been identified in other studies as having prognostic value for severe COVID-19 *(40, 41)*. Interestingly, these two genes also had significant (FDR ≤ 0.1) cis-eQTLs (**table S5**), indicating that there may be upstream genetic and epigenetic factors contributing to their differential expression in monocytes.

**Fig. 4.**
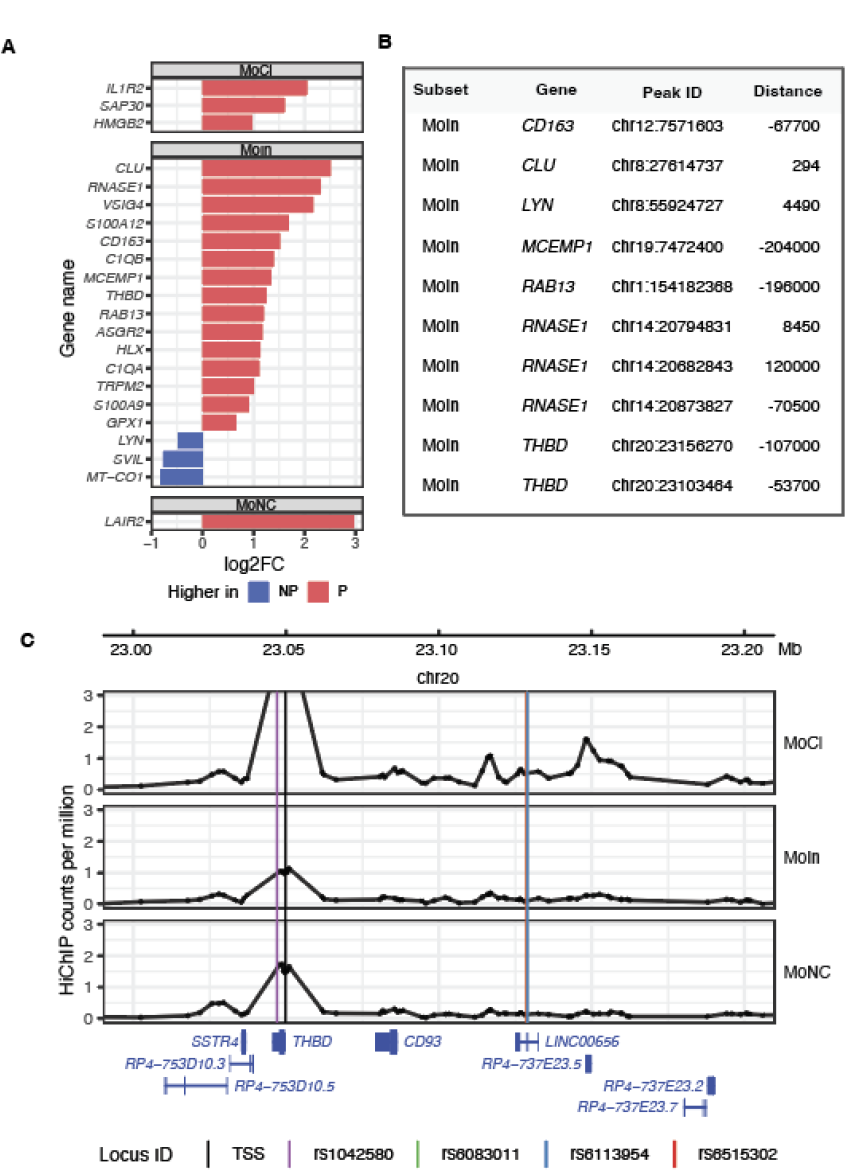
Monocyte Progressor versus Non-progressor DEGs. (**A**) Log2 fold-change (log2FC) of DEGs in classical monocytes (MoCl), intermediate monocytes (MoIn), and non-classical monocytes (MoNC) between Non-progressors (NP) and Progressors (P) at FDR ≤ 0.1. (**B**) DEGs with proximally associated DARs comparing all Progressor to Non-progressor cell subsets at FDR ≤ 0.1. Distance indicates the distance between the ATAC peak from the corresponding gene’s TSS. (**C**) Chromatin contact via HiChIP between regions with eQTL SNPs (rs6083011, rs6113954, and rs6515302) and the THBD TSS in monocyte subsets, with gene track and literature-reported SNP rs1042580 for context. The black vertical line represents the transcriptional start site (TSS), and colored vertical lines represent variant positions, with the lines for rs6083011, rs6113954, and rs6515302 overlapping in the figure.

Previous work identified a genetic variant in *THBD* (rs1042580) associated with risk of ICU admission and mortality from COVID-19 *(42, 43)*. This variant and the significant variants identified as eQTLs (rs6083011, rs6113954, rs6515302) were not independently associated with progression status in our study (unadjusted p > 0.4 for all four SNPs), which is not surprising given our relatively small study. However, chromatin contact mapping in healthy donors indicated that the 3 eQTL SNPs fall in regions with evidence of physical contact with the *THBD* promoter region in classical monocytes, but not intermediate monocytes (Fig. 4C **and fig. S7**). In contrast, rs1042580 is in the 3’UTR. Taken together, these results suggest that THBD may influence progression and identify two candidate regulatory mechanisms for *THBD*: 1) genetic variation in the three eQTL SNPs influencing chromatin contact and subsequent gene expression, and 2) genetic variation in the 3’UTR influencing another regulatory mechanism such as mRNA localization, stability, or translation *(44)*.

### Effector memory CD8^+^ T cells from Progressors show strong early activation and possible dysfunction

Several studies have linked strong activation in the CD8^+^ T cell compartment to severe COVID-19 *(16, 45, 46)*. Using GSEA, effector memory CD8^+^ T cells in Progressors showed a strong enrichment (p = 0.0073) of genes involved in an activation-related signature, displaying increased expression of markers of cell proliferation (*MKI67*), surface markers of activation (*CD38, HLA-DRA*), cytokines (*IFNG, TNF*), and cytolytic factors (*GZMA, GZMB, GNLY, PRF1, GZMK, GZMH*) (Fig. 5A). These changes were also apparent at the chromatin level, as Progressor effector memory CD8^+^ T cells showed increased chromatin accessibility around the cell proliferation marker Ki67 (*MKI67*) and granzyme (*GZMK, GZMH, GZMB, GZMA*) chromatin regions, as well as increased accessibility around activation-related regions (*CD38, HLA-DRA*) (Fig. 5B). In particular, granzyme A (*GZMA*) was increased in Progressors by differential analysis at FDR ≤ 0.1. These findings suggest that Progressors exhibit a strong activation signature in the effector memory CD8^+^ T cell compartment even at this early phase of infection.

**Fig. 5.**
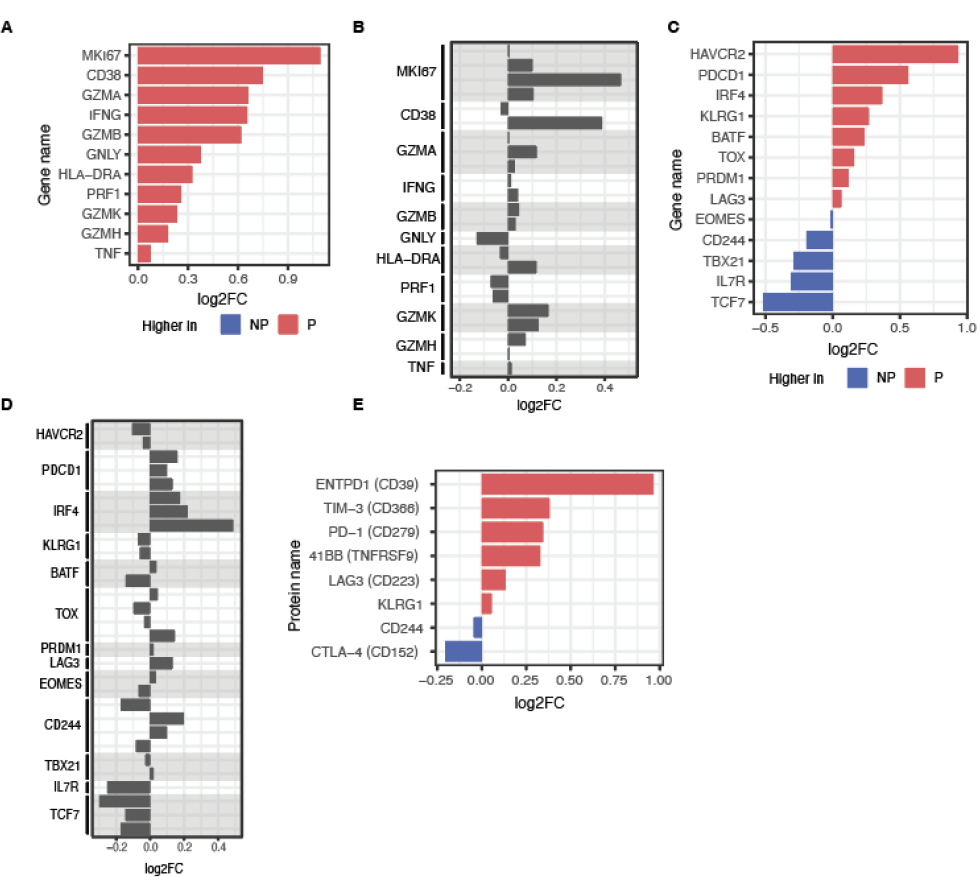
Highly activated phenotype in effector memory CD8+ T cells of Progressors. (**A**) Log2 fold-change (log2FC) of hyperactivation-related genes by RNA-seq in effector memory CD8^+^ T cells between Non-progressors (NP) and Progressors (P). (**B**) Log2FC in chromatin accessibility by ATAC-seq of hyperactivation-related genes in effector memory CD8^+^ T cells between Non-progressors and Progressors. Multiple bars indicate multiple identified peaks within +/− 1kb of gene TSS. (**C**) Log2FC of T cell dysfunction or activation-related genes by RNA-seq in effector memory CD8^+^ T cells between NP and P. (**D**) Log2FC in chromatin accessibility by ATAC-seq of dysfunction or activation-related genes in effector memory CD8^+^ T cells between Non-progressors and Progressors. Multiple bars indicate multiple identified peaks within +/− 1kb of gene TSS. (**E**) Log2FC of cell surface protein levels of regulatory and activation markers by TaPE-seq in effector memory CD8^+^ T cells between NP and P.

The effector memory CD8^+^ T cell subset also showed the highest number of DEGs (Fig. 2B). To better understand the function of these DEGs, we compared the 22 significant genes in Progressors (FDR ≤ 0.05) (**Fig. S8A**) with DEGs from a well characterized LCMV murine model of viral-induced T cell activation *(47–49)*. Twenty-one of these DEGS in Progressor effector memory C8^+^ T cells were differentially regulated in activated CD8^+^ effector T cells from LCMV-infected mouse spleens relative to naive splenic CD8^+^ T cells (**Fig. S8B**) *(50)*. Fourteen of the 21 genes matched the directionality of expression in the mouse model of T cell activation and the Progressor outcome group (i.e., increased in both Progressors and in the LCMV murine model, or decreased in both), suggesting that effector memory CD8^+^ T cells from Progressors may be enriched for genes involved in virally-induced T cell activation. Of the 144 DEGs detected at FDR ≤ 0.1, some effector-related functionality has been previously described for *LAMTOR1 (51)*, *ZNF395 (52)*, *ARPC1A (53, 54)*, *MECP2 (55)*, *CCL3 (56)*, and *GZMA (57)*.

Several studies have found indications of both effector- and dysfunction-related signatures in the CD8^+^ T cell compartment of SARS-CoV2-infected individuals *(46, 58, 59)*. We found that Progressor effector memory CD8^+^ T cells expressed increased levels of regulatory cell surface markers (*HAVCR2, PDCD1, KLRG1, LAG3*) *(60)* and effector-related transcription factors (*IRF4, BATF, TOX, PRDM1) (61, 62)* while simultaneously showing a decrease in the expression of memory or persistence-related markers (*TBX21, IL7R, TCF7*) *(63)* by GSEA (enrichment p = 0.081) (Fig. 5C), which could support either an effector or dysfunction-related signature. At the chromatin level, effector memory CD8^+^ T cells in Progressors showed increased accessibility at the *PDCD1* and *IRF4* regions while showing concurrent decreased accessibility around the *IL7R* region (Fig. 5D), providing some evidence of concordance at the chromatin level. Cell surface protein levels in Progressor effector memory CD8^+^ T cells also showed an increase in the levels of proteins related to regulatory receptors (TIM-3, PD-1, LAG3, KLRG1) and activation markers (CD39) (p = 0.004) (Fig. 5E). Altogether, these data support a model of strong CD8^+^ T cell activation, and possibly dysfunction, consistent with viral infection.

## DISCUSSION

Much of our current knowledge on the molecular pathogenesis of SARS-CoV-2 infection has been obtained at later phases of infection when patients have progressed in their disease. By enrolling patients soon after diagnosis and following them longitudinally, we were able to distinguish early differences in immune response between individuals on different courses of disease severity. Using multi-omic phenotyping of biospecimens collected at study enrollment, we identified a diverse range of molecular associations with COVID-19 progression.

Our findings highlight that cellular and molecular immunological differences occur early in disease between individuals who progress to clinically significant forms of COVID-19 and those who experience a milder disease trajectory. The differences observed between Progressors and Non-progressors manifest in blood proteins and circulating immune cells in both the myeloid and lymphocyte compartments. In general, the differences are consistent with an increased inflammatory or activated state in Progressors compared to Non-progressors.

One key observation of our study was the relatively higher abundance of multiple pro-inflammatory cytokines in Progressors, including prototypical mediators such as IL-6, IFNγ, and TNF. Cellular gene expression differences for *IL6* in monocytes and *IFNG* and *TNF* in T cells were directionally consistent with the plasma cytokine changes, although the expression changes did not reach significance and also might not reflect the biology of tissue-localized leukocytes. Many studies have reported a hyperinflammatory phenotype in hospitalized patients with severe COVID-19 *(64, 65)*, and interventional trials targeting inflammatory cytokines have shown some success in patients with severe disease *(23–25)*. Given the short time from diagnosis to molecular measurements in our study, the Progressors are unlikely to meet the criteria for the hyperinflammatory state described in severe COVID-19; rather, the molecular changes may reflect a precursor or incident hyperinflammatory state that could eventually evolve into hyperinflammation that meets clinical criteria for severe COVID-19. Importantly, this highlights that an incident hyperinflammatory state may be detected earlier than the onset of clinically severe disease.

Inflammatory cytokines such as IL-6, IFNγ, and TNF are also associated with myeloid and T cell-driven inflammatory states and autoimmune diseases. Notably, these are the cellular populations in which we see the largest differences between the Progressor and Non-progressor groups, and these are also the predominant cell types comprising lung infiltrates of patients with severe COVID-19 *(64)*. In monocytes, we observed a relative decrease in later-stage cellular populations (i.e., intermediate and non-classical monocytes) in the circulation, with expression changes in the intermediate monocyte population skewing towards a less mature, classical-like phenotype characterized by high levels of *CD163* expression. During the normal course of monocyte differentiation, classical monocytes evolve into intermediate and then nonclassical monocytes over the course of days, losing *CD163* expression, so one possible explanation for our findings is that classical monocytes are trafficking to sites of infection in the tissue and thus failing to differentiate into more mature monocyte subsets in the circulation. This possibility is consistent with the observation that CD163^+^ myeloid cells increase in lung tissue of individuals who have died from COVID-19 *(66)*. Of note, one of the genes we found to have increased expression in intermediate monocytes in Progressors, *S100A12*, was also previously identified as an amplifier of early immune responses in sepsis *(67)* and as having prognostic value for severe COVID-19 *(68)*.

There were two particularly interesting functional genomic signals from our study that are strongly supported by a broader set of literature. First, *THBD* was significantly different between Progressors and Non-progressors with regards to both DEGs and DARs in our work. A genetic variant in *THBD* (rs1042580) has been reported to confer genetic risk for ICU admission and mortality from COVID-19 *(42, 43)*, and we have identified additional candidate genetic risk factors with evidence of chromatin contact between the regions proximal to these risk factors and the promoter of *THBD. THBD* genetic risk factors are also associated with non-COVID venous thromboembolism *(69, 70)*, and thrombomodulin levels are associated with protection from ischemic stroke *(70, 71)*. Second, *MCEMP1*, which was also a significant DEG and DAR in our study, has been identified as a putative driver gene of critical COVID-19 in young patients *(72)* and as a prognostic molecule for severe COVID-19 *(40)* and ICU admission *(73)*. It has also been reported to be a key molecule associated with sepsis *(74)* and other acute inflammatory conditions *(73)*. Moreover, *MCEMP1* expression has prognostic value for stroke outcomes *(75)*, which points to a potentially overlapping mechanism (i.e., coagulopathy) with *THBD* in COVID-19 *(76, 77)*.

In the T cell compartment, we observed changes in the cytotoxic CD8^+^ T cells. Gene expression changes in these cells are largely consistent with a more highly activated state in Progressors, characterized by relative increases in molecular features associated with proliferation, cytokine production, and cytolytic functions. This included *IFNG* and *TNF*, whose protein products were notably also increased in the plasma of Progressors. This is only suggestive that CD8^+^ T cells could be the origin of these circulating pro-inflammatory cytokines, as other key candidate sources of cytokines such as tissue-infiltrating leukocytes were not interrogated, but the findings are consistent with other molecular associations with Progressors.

Importantly, these findings might suggest that there is a window of sub-clinical, “molecular progression” that precedes the clinical manifestations of COVID-19. Therefore, coupled with the right diagnostic tools, interventions could potentially be deployed earlier to those at highest risk of progressing to more severe disease. Some of the early signals reported herein could be a starting point for the development of diagnostic tools, although we have not evaluated the prognostic potential of the signals reported herein due to limitations of the study design. The detailed mechanistic findings also provide additional insight towards possible therapeutic approaches. Together, such tools could be important enablers of earlier management and reduction of severe COVID-19.

## MATERIALS AND METHODS

### Study design

The COVID Progression Retrospective (CPR) study is a retrospective cross-sectional study of existing datasets including Progressors and Non-progressors from two Verily-sponsored studies: COVID-19 Immune Response Study (Cove) and Predictors of Severe COVID-19 Outcomes (PRESCO). The CPR study was approved by Western Institutional Review Board (WIRB) (Protocol# 103678) on 8/30/2021. The Board found that this research meets the requirements for a waiver of consent under 45 CFR 46 116(f)[2018 Requirements] 45 CFR 46.116(d) [Pre-2018 Requirements].

*Cove.* Cove is a decentralized, prospective study collecting biological measurements and clinical and epidemiological data in participants confirmed positive for SARS-CoV-2 at the time of COVID testing, with the aim of characterizing molecular signatures associated with COVID-19 disease progression over 28 days. Adults testing positive for COVID-19 from community testing programs (Baseline COVID-19 Testing *(78)* and partner programs) and meeting the eligibility criteria were invited to enroll in the Cove study. Eligible participants were adults that 1) were 18 years old or older, 2) were U.S. residents, 3) tested positive for COVID-19 within the past 5 days, 4) were willing and able to provide informed consent, and 5) were willing and able to complete all study procedures. Additionally, participants were excluded if they 1) were pregnant or planning to become pregnant within the next 30 days, 2) had HIV infection or a history of cancer, 3) were undergoing treatment with immunosuppressants, 4) had known chronic or acute infections other than SARS-CoV-2, or 5) received any dose of a COVID-19 vaccine. A total of 115 participants were enrolled between January and May of 2021 across 5 states (NJ, CA, PA, TX, and NY).

Longitudinal clinical data and biospecimens were collected at up to 3 home visits and through daily electronic patient-reported outcomes (ePROs) following study enrollment, with home visits occurring on Days 3, 5, and 7 and an outcome survey on Day 28. The study was approved by WIRB (Protocol# 102857) at one decentralized site (Elligo Health Research). Written informed consent was obtained from all participants or their legally authorized representatives before study-related procedures were performed.

*PRESCO.* PRESCO (clinicaltrials.gov NCT04388813) is a multi-center, prospective, 3-month cohort study designed to identify clinical and molecular signatures associated with progression to severe COVID-19. Adults with laboratory-test confirmed acute SARS-CoV-2 infection (RT-PCR or antigen testing) and meeting the eligibility criteria were invited to participate. Eligible participants were adults that 1) were 18 years old or older, 2) were U.S. residents, 3) confirmed positive for COVID-19, 4) received care at a participating site (The University of Arizona, Cedars-Sinai Medical Center, University of Illinois Chicago, Rush University Medical Center, Weill Cornell Medical College, University of Texas Southwestern Medical Center, Baylor College of Medicine, and Inova Health Care Services), 5) were willing and able to provide informed consent, and 6) were willing and able to complete all study procedures. Participants were excluded if they were pregnant. A total of 494 patients were enrolled between May 2020 and June 2021.

Longitudinal clinical data and biospecimens were collected at up to five visits during SARS-CoV-2 infection and recovery: (1) enrollment during initial hospital presentation, and if occurred, (2) two days after hospitalization; (3) on the day of admission to intensive care unit (ICU); (4) the day of hospital discharge; and (5) approximately 3 months after hospital presentation. The study was approved by a central WIRB (Protocol# 20201016) and at each of the participating sites. Written informed consent was obtained from all participants or their legally authorized representatives before study-related procedures were performed.

*CPR.* All datasets from participants enrolled in the Cove study were eligible to be included in CPR. For a participant dataset from PRESCO to be included, the participant must have tested positive for SARS-CoV-2 within 5 days prior to or after enrollment in the PRESCO study. Additionally, datasets from PRESCO participants that met the following criteria were excluded from the CPR study: 1) had HIV infection or a history of cancer, 2) dexamethasone treatment before enrollment, 3) any immunosuppressive therapy within the prior 14 days, 4) known chronic or acute infection other than SARS-CoV-2, and 5) received any dose of COVID-19 vaccine. In total, 83 participant datasets from PRESCO were combined with the 115 from Cove, for a total of 198 participant datasets eligible for inclusion in the CPR study.

### Assessment of COVID-19 progression

#### Protocol definition

Disease progression was defined as hospitalization or outpatient treatment ‘related to’ COVID-19 worsening or the worsening of a pre-existing condition triggered by SARS-CoV-2 infection within 28 days. Outpatient treatment definition excluded any over-the-counter (OTC) medications such as antipyretics, antitussives, and analgesics, but included treatments such as supplemental oxygen, intravenous fluids, diuretics, anticoagulation, increase in dose of medications for pre-existing condition(s), and antibiotics.

#### Challenges to implementation

The definition of progression was reliant on ‘relatedness’ of the treatment to the SARS-CoV-2 infection, with ‘relatedness’ intended to act as an indication of progression of clinical symptoms. A number of challenges existed to a simple, algorithm-based approach to labeling Progressors. Both Cove and PRESCO studies were conducted during the height of the pandemic amidst rapidly evolving knowledge and treatment recommendations. In addition, in some cases, the reason for hospitalization was nuanced. For example, in PRESCO, where patients were enrolled upon presentation at the hospital, a positive SARS-CoV-2 test may have been an incidental finding and not the reason for the visit (e.g., among reasons for hospitalization listed: ‘suicidality’ and ‘sickle cell anemia’). In either study, patients may have been briefly hospitalized and provided with supplemental oxygen in the absence of any relevant clinical symptoms, e.g., shortness of breath or low blood oxygenation, out of a presumed abundance of caution due to advanced age or serious underlying comorbid condition. Misclassification of Progressors was seen as a very serious risk to addressing the CPR study’s objectives.

#### Adjudication process

In light of the treatment landscape and the nuanced clinical data, as well as the differing data collection between the two studies, clinical expertise was sought to adjudicate the progression outcomes. An adjudication panel was convened, consisting of 3 Janssen clinicians with experience and expertise in respiratory viral infections and the conduct of clinical trials and who were not involved in this study. Per the adjudication charter, panel members reviewed full casebooks for all patients eligible to be included in the CPR study, in order to determine whether or not they met the definition of a Progressor. Each member provided an independent Progressor ‘vote’ for each patient; non-unanimous decisions were discussed and debated, with a two-thirds majority vote being sufficient to finalize the progression decision. Non-progressor patients represented individuals in the study that did not meet the progression criteria within 28 days. In some cases where the adjudication panel did not have enough information, no progression outcome was assigned, and the participant status was labeled as undetermined.

### Multi-omic analysis

All comparisons within this work were between Progressors and Non-progressors, unless otherwise specified.

Verily’s Immune Profiler platform was employed to conduct multi-omic analysis. This platform begins with the isolation of 25 immune cell subsets (5 myeloid cell subsets, 7 B cell subsets, 10 T cell subsets, 2 NK cell subsets, and a bulk PBMC sample) from a starting material of approximately 10 million cryopreserved PBMCs per individual. ATAC-seq and RNA-seq are measured for each of the 25 subsets, and TaPE-seq is performed for the 12 subsets in the T and NK panel (**table S3**). Additionally, WGS data were generated as permitted by participant consent.

#### FACS

Frozen cryovials of PBMCs in liquid nitrogen were thawed in a 37 °C water bath and transferred to a 1.5 mL Eppendorf tube. 500 μL of warmed R10 media was added to the 1.5 mL tube and left for 2 minutes to come to equilibrium. The cells were then centrifuged for 5 minutes at 500 × *g*, 25 °C. The cell pellet was resuspended with 1 mL of warm FACS buffer, and the cells were counted. The cells were then centrifuged again for 5 minutes at 500 x *g*, 4°C. The cell pellet was resuspended in 50 μL of FACS buffer with 5 μl of BD Human Fc Block and incubated for 5 minutes. 50 μL of staining cocktail was added per 10 million cells counted for the respective flow cytometry panels to be analyzed (T cell, B cell, myeloid panel) and incubated for 15 minutes at 4 °C in the dark. Cells for B cell panels and myeloid panels were washed in FACS buffer, resuspended in a final volume of 500 μL FACS buffer, and passed through a 35 μm cell strainer cap. 5 μL of 7-AAD live/dead dye was added to the cells before sorting. Cells for the T cell panels were washed in FACS buffer, stained with 250 µL of OligoAb cocktail (Supplementary Materials and Methods), incubated for an additional 15 minutes at 4 °C in the dark, re-washed in FACS buffer, resuspended in a final volume of 500 μL FACS buffer, and passed through a 35-μm cell strainer cap. 5 uL of 7AAD live dead dye was added to the cells before sorting. Stained samples were sorted on a FACSAria Fusion (BD Biosciences, San Jose, CA). Using FACSDiva v8.0.1 software, the samples were gated first by forward and side scatter properties, then FSC-H vs FSC-A for singlet discrimination, and finally, with their respective markers for each cell type (**table S3**). For each cell type of interest, 800-10,000 cells per sample were sorted into the tagmentation buffer. Additionally, a minimum of 500,000 cells were recovered from either the PBMCs or a separate buffy coat aliquot for whole genome sequencing.

#### Hybridization and Separation

RNA was separated from the other components, including tagmented DNA and antigens bound by barcoded antibodies in the cell-containing samples, for further analysis. Biotin-OligodTVN (in house preparation) beads were added to each sample, mixed, and beads captured using a magnet. The supernatant was aspirated and the plate was removed from the magnet. The beads were resuspended with the lysed cells for 30 minutes at 25 °C with orbital mixing at 1500 rpm using an Eppendorf Thermomixer (Eppendorf, Hamburg, Germany). Samples were centrifuged and placed on a magnet, and the supernatant was transferred into a new plate for subsequent ATAC-seq and TaPE-seq processing. The plate containing the RNA samples was immediately processed through the RNA-seq workflow.

*ATAC-seq and TaPE-seq*.

ATAC-seq was performed as previously described *(79)*, with the exception that barcoded oligos from proteins bound by the OligoAb cocktail (Supplementary Materials and Methods) were simultaneously captured in the supernatant during the above hybridization, and a separation step was conducted for protein estimation using TaPE-seq *(80, 81)*.

#### RNA-seq

RNA-seq was performed using a SMART-Seq2-based procedure optimized for use on hundreds of cells *(82)*.

#### Library quality control, pooling, and sequencing

RNA-seq libraries and libraries containing both ATAC-seq and TaPE-seq were quantified by qPCR using the KAPA Library Quantification Kit (Complete kit, Universal) (F. Hoffmann-La Roche AG, Basel, Switzerland) on the CFX384 Touch™ Real-Time PCR Detection System (Bio-Rad Laboratories Inc, Hercules, CA, USA). Libraries were pooled and sequenced utilizing a two-pass approach: pools of ATAC-seq and TaPE-seq libraries and pools of RNA-seq libraries were pooled in equimolar ratios and sequenced across a single lane of a flow cell each; libraries were re-pooled after this first-pass sequencing accounting for both their qPCR quantification and the number of first-pass sequencing reads achieved to obtain a more even read distribution across the pool during second-pass sequencing. Libraries were sequenced to total target depths of 10,000,000 reads for RNA-seq, 20,000,000 reads for ATAC-seq, and 50,000 reads for TaPE-seq on the Illumina NovaSeq 6000 platform (Illumina Inc, San Diego, CA, USA), with v1.5 chemistry kits generating paired-end (2×150 bp) reads.

#### Whole Genome Sequencing

WGS was performed on isolated cells using a PCR-free procedure. Briefly, a total of 350 ng of gDNA from each sample based on Quant-iT Picogreen quantification (Thermo Fisher Scientific, Waltham, MA, USA) was mechanically fragmented to a target size between 350 and 400 base pairs (bp) on the Covaris LE220 focused ultrasonicator (Covaris, Woburn, MA, USA). The sheared gDNA underwent size selection using AMPure XP beads (Beckman Coulter, Brea, CA, USA) to tighten the size distribution of the gDNA fragments. The size selected gDNA was end repaired, A-tailed, and adapter-ligated using the KAPA PCR-free Hyper Prep Kit in combination with KAPA Unique Dual-Indexed Adapters (F. Hoffmann-La Roche AG, Basel, Switzerland). Adapter-ligated libraries were purified by AMPure XP bead cleanup. Library yields were assessed by qPCR using the KAPA Library Quantification Kit (Complete kit, Universal) (F. Hoffmann-La Roche AG, Basel, Switzerland) on the CFX384 Touch™ Real-Time PCR Detection System (Bio-Rad Laboratories Inc, Hercules, CA, USA). Dual-indexed libraries were subsequently pooled and sequenced to a target depth of 30X coverage on the Illumina NovaSeq 6000 platform (Illumina Inc, San Diego, CA, USA), with v1.5 chemistry kits generating paired-end (2×150 bp) reads.

### Chromatin contact maps

#### HiChIP

Frozen cryovials of 100 million PBMCs (STEMCELL Technologies, Vancouver, BC, Canada) were sorted by FACS following the method described above for three donors. For each cell type of interest, a target of minimally 50,000 cells were collected and crosslinked with 1% formaldehyde for 10 minutes at room temperature, quenched with 125 mM glycine for 5 minutes at room temperature, and then pelleted and washed in preparation for HiChIP library preparation. *In situ* contact libraries were generated according to the *in situ* HiChIP published protocol *(83)* with the following optimizations: sonication was performed in Covaris milliTubes at the following settings 300 PIP, 15% DF, and 200 CPB for 4 minutes, clarification was reduced to 14,000 rcf for 10 minutes, samples were washed following immunoprecipitation twice in Low Salt Wash Buffer and twice in High Salt Wash Buffer (no LiCl washes were completed) and final library size selection utilized SPRISelect Beads (Beckman Coulter, Indianapolis, IN, USA). A cutoff of 0.15 counts per million was applied to determine the presence of chromatin contact. This cutoff was optimized for enrichment of significant eQTLs and represents the 95th percentile of quantified HiChIP loops.

### Quantification of viral load

Viral load was quantified using droplet digital PCR (ddPCR) from mid-turbinate swabs collected at each visit. Extraction of SARS-CoV-2 RNA was performed using the MagMax Viral/Pathogen Nucleic Acid Isolation kit (Thermo Fisher Scientific, Waltham, MA) following manufacturer’s instructions, and ddPCR was performed using a previously described protocol *(84)*. In addition to quantifying the *N* and *ORF1a* SARS-CoV-2 genes, *RPP30* was quantified as a positive control for swabbing using publicly available sequences *(85)*.

### Quantification of plasma cytokines

Forty-seven cytokines (sCD40L, EGF, Eotaxin, FGF-2, Flt-3 ligand, Fractalkine, G-CSF, GM-CSF, GROα, IFNα2, IFNγ, IL-1α, IL-1β, IL-1ra, IL-2, IL-3, IL-4, IL-5, IL-6, IL-7, IL-8, IL-9, IL-10, IL-12p40, IL-12p70, IL-13, IL-15, IL-17A, IL-17E/IL-25, IL-17F, IL-18, IL-22, IL-27, IP-10, MCP-1, MCP-3, M-CSF, MDC, MIG, MIP-1α, MIP-1β, PDGF-AA, PDGF-AB/BB, TGFα, TNF (TNFα), LTA (TNFβ), and VEGF-A) were quantified from plasma samples using the MILLIPLEX MAP Human Cytokine/Chemokine/Growth Factor Panel A on a Luminex FLEXMAP 3D instrument according to manufacturer’s instructions. For each cytokine measured, values lying outside of the standard curve were imputed to the nearest standard concentration. Three cytokines (GM-CSF, IL-17F, and IL-3) were excluded from further analysis because >80% of their measurements fell outside of their respective standard curves. Forty-four cytokines were thus used for downstream analysis. Individual cytokine measurements that did not have either a) bead counts ≥35 and technical CV ≤30%, or b) bead counts ≥20 and technical CV ≤15%, were also excluded from analysis.

### Differential Analysis

To better isolate the relationship between molecular features and progression, a number of covariates are adjusted for in our models unless otherwise stated. These covariates include clinical and demographic variables, assay-specific variables, and a variable to adjust for the study from which a biospecimen was collected. Clinical and demographic variables were individually assessed using logistic regression to determine each variable’s unadjusted association with progression. All variables that met the criterion of p ≤ 0.20 were considered for inclusion in the “full model” - a multivariable logistic regression with progression as the outcome variable *(86)*. Beginning with the full model, variables with a p-value > 0.10 were removed in a stepwise fashion in order of descending p-value. Age and sex were considered key covariates and were thus never removed from the model. The full model also included race, body mass index, and indicators for the following comorbid conditions: cardiovascular and cerebrovascular, metabolic and/or diabetic, renal, and respiratory. After variable selection, the clinical covariates retained in the final model included age, sex, race, and an indicator for comorbid respiratory conditions. Association analyses with Immune Profiler data additionally adjusted for cell viability, neutrophil frequency (as a measure of neutrophil contamination during PBMC isolation), and immune subset recovery count, as we have found these variables to be associated with sample quality. Lastly, principal component analysis of the Immune Profiler data showed a batch effect associated with the study from which biospecimens were collected. Therefore we included an indicator variable to adjust for study.

Univariate differential analysis to identify molecular features differentiating Progressors from Non-progressors was performed using linear modeling methods. For these analyses, each molecular feature was regressed on the outcome group and appropriate clinical and technical covariates. For count-based data such as RNA-, ATAC-, and TaPE-seq, the voom-limma method was used *(87)*. The method normalizes data using the default “TMM” method in the edgeR package, estimates the mean-variance relationship of the log-counts in order to generate sample weights, and conducts statistical inference of the estimate of association with limma’s empirical Bayes analysis pipeline. For non-count-based data (e.g., cell subset frequencies and cytokine levels), differential analysis was performed by fitting generalized linear models (GLMs). Where appropriate, data were transformed (e.g., log transformation) prior to fitting the GLMs.

Following linear modeling, the Benjamini-Hochberg procedure was used to correct for multiple hypothesis testing within each molecular data type. For Immune Profiler data, comparisons were done per cell subset, and the resulting p-values across all tests within a cell subset were corrected for. Significance was assessed at FDR ≤ 0.05; but due to the small study size and potential loss of biological signal due to adjustment for multiple covariates, we occasionally looked more broadly for differential features at FDR ≤ 0.1.

### Pathway Analysis

Gene sets from MSigDB were used for pathway analysis, and three independent methods were employed. First, enrichment of gene sets for significant differential genes were tested using hypergeometric tests. Second, GSEA *(88)* was performed using effect estimates from univariate differential analysis, and enabled identification of gene sets where the individual genes may not be significantly differentially expressed but are nevertheless coordinated in their association to progression status. And third, pathway analysis using GSVA was done, and additional differential analysis of the GSVA pathway enrichment scores was performed *(89)*.

## Supplementary Materials

Supplementary Tables and Figures

Supplementary Materials and Methods

## Acknowledgements

We thank the following individuals/groups for their involvement: the Adjudication Panel - Lorant Leopold, Rekha Sinha, and Erika Van Landuyt - at Janssen Research and Development, LLC for their provision of adjudicated progression outcome as described in the methods sections; Charlie Knights at Janssen Research and Development, LLC for establishing the collaboration and securing financing; Inge Verbrugge at Janssen Research and Development, LLC for various data analysis and results discussions; the PRESCO study staff for acting as site PIs, investigators, project managers, coordinators, data managers, research staff, regulatory staff, and budget coordinators; the following individuals/teams at Verily Life Sciences, LLC for their contributions: the Clinical Data Management and Clinical Operations teams for coordinating sites and managing clinical data; the Member and Patient Services team for technical and patient support; the Molecular Production team for sample processing; the Sponsor Operations team for overall study program management; Sarah Chen, MD, for managing project governance and relationship alliance; Taylor Oniskey for biospecimen logistics management and molecular study management; Susan Kram Sousa for project management and relationship alliance; Hannah Polikowsky, PhD, for project management. Finally, thank you to all of the individuals who consented to participate in our study and generously provide their data and biological specimens. Funding for the study was provided by Verily and Janssen. KAD, DT, SV, VK, VT, JM, CCK, JAK, MS, MB, IOR, CRD, VFT, SP contributed to conceptualization; KAD, DT, GT, SV, SK, VK, MJW, SAS, PK, VT, MVL, JM, CCK, JAK, CC contributed to methodology; GT, MJW, AMS, SAS, PK, CC contributed to software; DT, GT, VK, MJW, AMS, SAS, PK, VT, MB, CC contributed to validation; GT, SK, VK, MJW, AMS, VT, MVL contributed to formal analysis; JTL, JML, BY, MRM, KMF, VT, MS, MB, IOR, CRD, JNM, VFT, SP contributed to investigation; JM, CCK, MB, JNM, VFT, SP contributed to resources; DT, GT, SV, SAS, PK, VT, MVL, IOR, JNM, CC contributed to data curation; KAD, GT, JTL, SK, JML, BY, KMF, CCK contributed to writing (original draft prep); KAD, DT, GT, SV, VK, JML, BY, JTL, SK, SAS, MRM, KMF, VT, MVL, JM, CCK, JAK, MS, MB, IOR, CRD, JNM, VFT, SP contributed to writing (review+ editing); GT, JTL, SK, JML, BY, MJW, AMS contributed to visualization; KAD, DT, GT, JTL, SV, VT, MVL, JM, CCK, JNM contributed to supervision; KAD, DT, GT, VK, CCK contributed to project administration; DT, JM, JNM contributed to funding acquisition.DT, SV, VK, VT, MVL are employed by Janssen Research & Development, LLC. KAD, GT, JTL, SK, JML, BY, MJW, SAS, PK, KMF, CCK, CC maintain equity ownership and employment at Verily Life Sciences LLC, an Alphabet subsidiary. KAD, GT, JTL, SK, JML, BY, CCK are inventors listed on U.S. Application No. 63/496,918. SP reports personal fees from Jazz Pharmaceuticals, Inc., other from UpToDate, Inc., grants from NIH (R25-HL126140, R33-HL151254; OT2-HL161847; R21-HD109777; C06-OD028307; HL140144; HL138377; OT2-HL156912 and OT2HL158287), grants from PCORI (DI-2018C2-13161, CER-2018C2-13262), grants from Department of Defense (W81XWH20C0051 and W81XWH2110025), grants from Pima County Health Department (CPIMP211275), grants from Arizona Commerce Authority (LTR DTD 021822), grants from Philips, Inc. (0483-06-161311-73077), grants from Sommetrics, Inc., grants from American Academy of Sleep Medicine Foundation (AASMF; 169-SR-17), grants from Regeneron, Inc.; SP also has a patent US20160213879A1 licensed to SaiOx, Inc. CRD serves on advisory boards for Abbott Diagnostics, Ortho/Quidel Diagnostics, and Roche Diagnostics. CRD is an inventor of a method for assessing differential risk for developing heart failure (patent number 10509044). JAK is funded by the U.S. National Institutes of Health / National Heart, Lung, and Blood Institute, COPD Foundation, Regeneron, U.S. Patient Centered Outcomes Research Institute, American Lung Association; JAK has also provided consulting for GlaxoSmithKline, AstraZeneca, CereVu Medical, Propeller/ResMed, BData, Inc. A limited set of data are available upon request for the purpose of quality assessment.

