## Supplemental tables, figures, and methods for "Multi-omic Profiling Reveals Early Immunological Indicators for Identifying COVID-19 Progressors"

### SUPPLEMENTARY TABLES AND FIGURES

**Table S1. Characteristics of enrolled CPR study participants.**

| Characteristics | Total (N = 162) | Progressor (N = 24) | Non-progressor (N = 138) |
|---|---|---|---|
| BMI, mean (SD) | 29.6 (7.4) | 32 (7) | 29 (7) |
| BMI category*, N (%) | | | |
| Underweight | 1 (0.6%) | 0 (0%) | 1 (0.7%) |
| Healthy | 43 (27.2%) | 5 (20.8%) | 38 (28.4%) |
| Overweight | 54 (34.2%) | 7 (29.2%) | 47 (35.1%) |
| Obese | 45 (28.5%) | 7 (29.2%) | 38 (28.4%) |
| Severely obese | 15 (9.5%) | 5 (20.8%) | 10 (7.5%) |
| BMI missing, N (%) | 4 (2.5%) | 0 (0%) | 4 (3%) |
| Current smoker, N (%) | 42 (25.9%) | 7 (29.2%) | 35 (25.4%) |
| Smoking frequency#, N (%) | | | |
| Light use (<1-5 times per day) | 11 (26.2%) | 1 (14.3%) | 10 (28.6%) |
| Moderate use (6-10 times per day) | 10 (23.8%) | 1 (14.3%) | 9 (25.7%) |
| Heavy use (11+ times per day) | 2 (4.8%) | 0 (0%) | 2 (5.7%) |
| Missing | 19 (45.2%) | 5 (71.4%) | 14 (40%) |
| Years smoked#, N (%) | | | |
| <5 years | 9 (21.4%) | 1 (14.3%) | 8 (22.9%) |
| 5-15 years | 7 (16.7%) | 0 (0%) | 7 (20%) |
| 15+ years | 17 (40.5%) | 5 (71.4%) | 12 (34.3%) |
| Missing | 9 (21.4%) | 1 (14.3%) | 8 (22.9%) |
| Comorbid conditions, N (%) | | | |
| Autoimmune | 1 (0.6%) | 1 (4.2%) | 0 (0%) |
| Cardiovascular / cerebrovascular | 37 (22.8%) | 12 (50%) | 25 (18.1%) |
| Hematologic | 6 (3.7%) | 1 (4.2%) | 5 (3.6%) |
| Metabolic | 25 (15.4%) | 8 (33.3%) | 17 (12.3%) |
| Neurologic | 4 (2.5%) | 1 (4.2%) | 3 (2.2%) |
| Renal | 7 (4.3%) | 3 (12.5%) | 4 (2.9%) |
| Respiratory | 16 (9.9%) | 5 (20.8%) | 11 (8%) |
| Other | 49 (30.2%) | 8 (33.3%) | 41 (29.7%) |

** Percentages calculated out of participants with non-missing data*
*# Percentages calculated out of current smokers*

**Table S2. Clinical predictors of progression.**

| Characteristic | Category | Coefficient | p-value |
|---|---|---|---|
| **Age (continuous)** | **per year** | 0.059514 | **<0.001** |
| **Age (categorical)** | 18-29 (ref) | – | – |
| | **30-49** | 1.098612 | **0.17** |
| | **50-64** | 1.394878 | **0.10** |
| | **65+** | 3.152736 | **<0.001** |
| **Age (binary)** | **65+ vs. not** | 2.182299 | **<0.001** |
| **Sex** | **Female (vs. male)** | 1.347074 | **<0.05** |
| **Race** | **White (vs. not)** | -0.985058 | **<0.05** |
| | **Black (vs. not)** | 1.364757 | **<0.01** |
| Ethnicity | Hispanic (vs. not) | -0.236389 | 0.66 |
| Smoking | Current (vs. not) | 0.192078 | 0.69 |
| **BMI** | **per unit** | 0.051757 | **0.052** |
| BMI (categorical) | Under / Healthy (BMI <25) (ref) | – | – |
| | Overweight (BMI 25-<30) | 0.149886 | 0.81 |
| | Obese (BMI 30-<40) | 0.362448 | 0.56 |
| | **Severely Obese (BMI 40+)** | 1.360977 | **0.060** |
| **BMI (binary)** | **Severely Obese (vs. not)** | 1.182695 | **<0.05** |
| **Comorbidities** | Autoimmune (vs. not) | 22.357828 | 1.00 |
| | **Cardio/Cerebrovascular (vs. not)** | 1.508512 | **<0.01** |
| | Hematologic (vs. not) | 0.145417 | 0.90 |
| | **Metabolic (vs. not)** | 1.269430 | **<0.05** |
| | **Metabolic, Diabetes (vs. not)** | 1.756041 | **<0.01** |
| | **Metabolic, Other (vs. not)** | 0.928363 | **0.11** |
| | Neurologic (vs. not) | 0.671168 | 0.57 |
| | **Renal (vs. not)** | 1.565635 | **0.050** |
| | **Respiratory (vs. not)** | 1.111291 | **0.061** |
| | Other (vs. not) | 0.167992 | 0.72 |

**Table S3. Immune cell types profiled.**

| | | | | | |
|---|---|---|---|---|---|
| PBMCs | Myeloid | Dendritic cells | **coDC** | Conventional DCs | CD3- CD19- CD20- CD56- CD15- HLA-DR+ CD14- CD16- CD11c+ CD123- |
| | | | **plDC** | Plasmacytoid DCs | CD3- CD19- CD20- CD56- CD15- HLA-DR+ CD14- CD16- CD11c- CD123+ |
| | | Monocytes | **MoCl** | Classical monocytes | CD3- CD19- CD20- CD56- CD15- HLA-DR+ CD33+ CD91+ CD14++ CD16- |
| | | | **MoIn** | Intermediate monocytes | CD3- CD19- CD20- CD56- CD15- HLA-DR+ CD33+ CD91+ CD14++ CD16+ |
| | | | **MoNC** | Nonclassical monocytes | CD3- CD19- CD20- CD56- CD15- HLA-DR+ CD33+ CD91+ CD14- CD16+ |
| | T cells | CD4+ αβ T cells | **T4nv** | Naive CD4+ T cells | CD19- CD14- CD56- CD91- gadT- CD3+ CD4+ CCR7+ CD45RA+ |
| | | | **T4cm** | Central memory CD4+ T cells | CD19- CD14- CD56- CD91- gadT- CD3+ CD4+ CCR7+ CD45RA- |
| | | | **T4em** | Effector memory CD4+ T cells | CD19- CD14- CD56- CD91- gadT- CD3+ CD4+ CCR7- CD45RA- |
| | | | **T4ra** | CD45RA+ effector memory CD4+ T cells | CD19- CD14- CD56- CD91- gadT- CD3+ CD4+ CCR7- CD45RA+ |
| | | | **Treg** | CD4+ T regulatory cells | CD19- CD14- CD56- CD91- gadT- CD3+ CD4+ CD127- CD25+ |
| | | CD8+ αβ T cells | **T8nv** | Naive CD8+ T cells | CD19- CD14- CD56- CD91- gadT- CD3+ CD8+ CCR7+ CD45RA+ |
| | | | **T8cm** | Central memory CD8+ T cells | CD19- CD14- CD56- CD91- gadT- CD3+ CD8+ CCR7+ CD45RA- |
| | | | **T8em** | Effector memory CD8+ T cells | CD19- CD14- CD56- CD91- gadT- CD3+ CD8+ CCR7- CD45RA- |
| | | | **T8ra** | CD45RA+ effector memory CD8+ T cells | CD19- CD14- CD56- CD91- gadT- CD3+ CD8+ CCR7- CD45RA+ |
| | | γδ T cells | **gadT** | Gammadelta T cells | CD19- CD14- CD56- CD91- CD3+ gdTCR+ |
| | NK | NK cells | **NKhi** | CD56hi NK cells | CD3- CD19- CD14- CD91- CD56++ |
| | | | **NKlo** | CD56low NK cells | CD3- CD19- CD14- CD91- CD56+ |
| | B cells | Naive B cells | **BnUS** | Unswitched naive B cells | CD3- CD14- CD19+ CD10- CD38- CD20+ CD27- CD21+ IgM+ IgD+ |
| | | | **BnCS** | Class switched naive B cells | CD3- CD14- CD19+ CD10- CD38- CD20+ CD27- CD21+ IgM- IgD- |
| | | | **traB** | Transitional B cells | CD3- CD14- CD19+ CD10+ CD20+ CD38+ CD21+ CD24+ IgD+ |
| | | Effector B cells | **cMBc** | Class switched classical memory B cells | CD3- CD14- CD19+ CD10- CD38- CD20+ CD27+ CD21+ IgM- IgD- |
| | | | **MBim** | IgM+ IgD- classical memory B cells | CD3- CD14- CD19+ CD10- CD38- CD20+ CD27+ CD21+ IgM+ IgD- |
| | | | **aMBc** | Atypical Memory B cells | CD3- CD14- CD19+ CD10- CD38- CD20+ CD27- CD21- IgM- IgD- |
| | | | **pBcs** | Class switched plasmablast | CD3- CD14- CD19+ CD10- CD38++ CD20- IgM- IgD- |

**Table S4. Assay features measured.**

| Molecular assays | Features measured |
|---|---|
| Viral load | 3 |
| Luminex - cytokine panel | 47 |
| Immune Profiler | |
| Cell frequencies | 74 |
| RNA-seq | 60,049 (x24 subsets) |
| ATAC-seq | 260,056 (x24 subsets) |
| TaPE-seq | 117 (x24 subsets) |

Fig. S1.

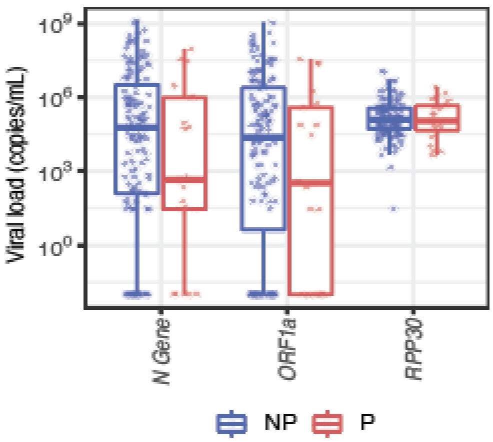


**Fig. S1. Viral load from mid-turbinate swabs.** Viral load in copies/ml for Non-progressors (NP) versus Progressors (P) for the N and ORF1a SARS-CoV-2 genes and the positive control gene, RPP30.

**Fig. S2.**

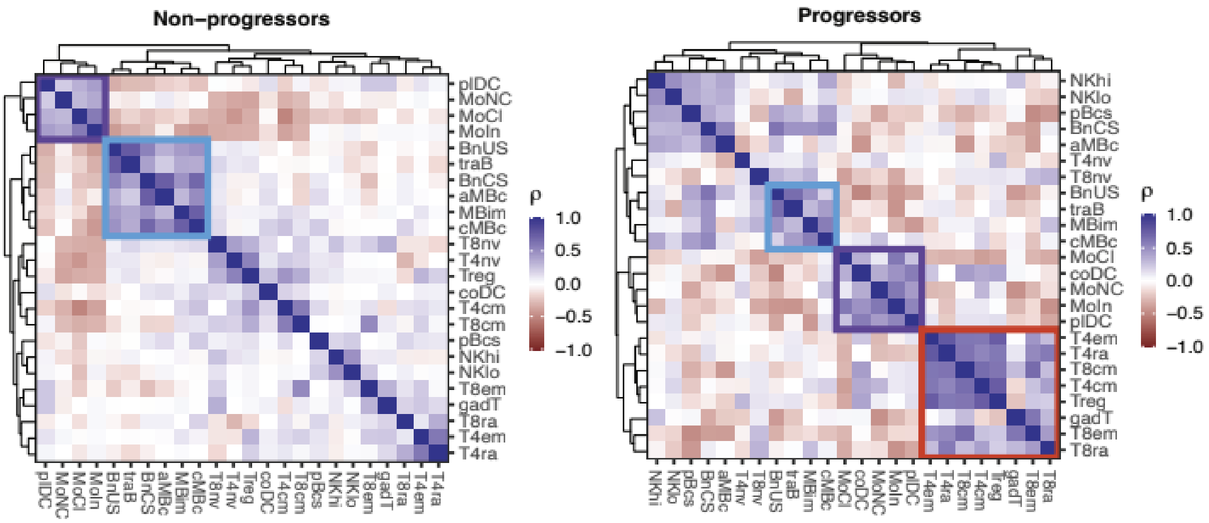


**Fig. S2. Cell subset frequency correlation heatmaps.** Heatmaps show pairwise Spearman's correlations within Non-progressor (left) and Progressor (right) cohorts. Blue, purple, and orange boxes indicate correlated clusters of B cell, myeloid cell, and effector T cell subsets, respectively; Abbreviations for cell subsets are defined in table S3.

**Fig. S3.**

A

B

C

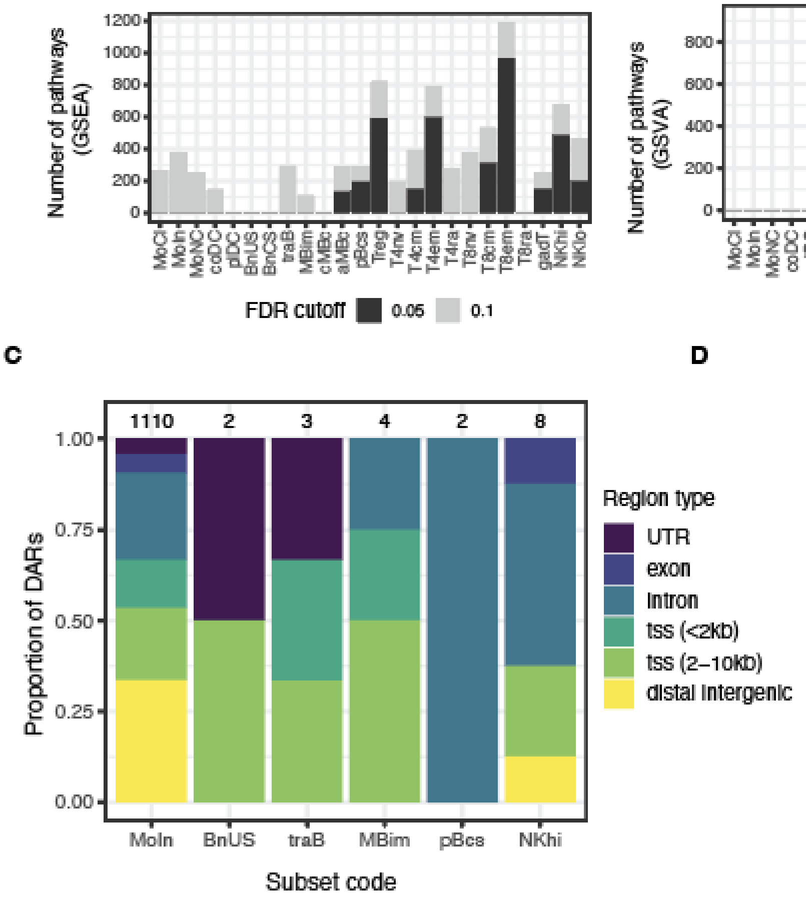


D

| subset code | LOOCV AUC |
|---|---|
| MoIn | 0.845 |
| T4nv | 0.774 |
| T8em | 0.709 |
| BnUS | 0.708 |
| cMBc | 0.699 |
| aMBc | 0.692 |
| pBcs | 0.681 |
| NKlo | 0.657 |
| T8nv | 0.650 |
| T4cm | 0.638 |
| coDC | 0.637 |
| MoNC | 0.635 |
| MoCl | 0.634 |
| NKhi | 0.632 |
| T8cm | 0.624 |
| BnCS | 0.622 |
| Treg | 0.611 |
| T8ra | 0.603 |
| plDC | 0.601 |
| traB | 0.592 |
| MBim | 0.576 |
| gadT | 0.569 |

**Fig. S3. Molecular pathway analysis and ML modeling.** (**A**) Identified number of significant pathways by gene set enrichment analysis (GSEA) pathway analysis between Non-progressors and Progressors by cell subset from RNA-seq data. (**B**) Identified number of significant pathways by gene set variation analysis (GSVA) pathway analysis between Non-progressors and Progressors by cell subset from RNA-seq data. (**C**) The proportion of DARs at FDR ≤ 0.1 within each subset found within each annotated genomic region. The total number of DARs discovered is indicated above each bar. (**D**) Block sparse partial least squares-discriminant analysis performed for each cell subset using RNA-seq, ATAC-seq, and progression outcome data assessed the classification performance for each subset. Cell subsets were ranked in their performance by the area under the ROC curve generated from Leave-one-out cross-validation (LOOCV); Abbreviations for cell subsets are defined in table S3.

**Fig. S4.**

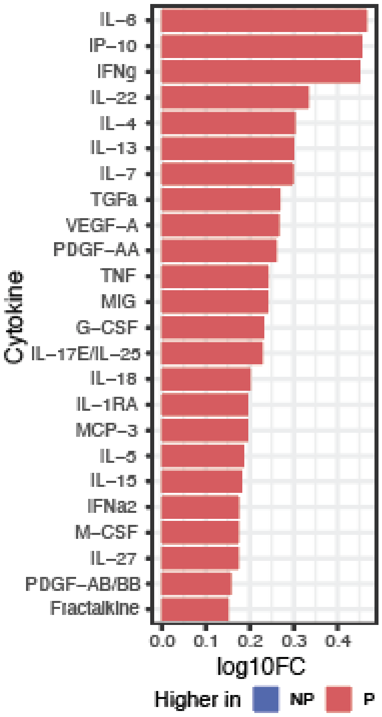


**Fig. S4. List of plasma cytokines and chemokines elevated in Progressors at FDR ≤ 0.1.** Cytokines and chemokines elevated in plasma of Non-progressors (NP) versus Progressors (P) at a FDR ≤ 0.1 as measured by Luminex assay. Comparative values are expressed as log10 fold change (log10FC).

**Fig. S5.**

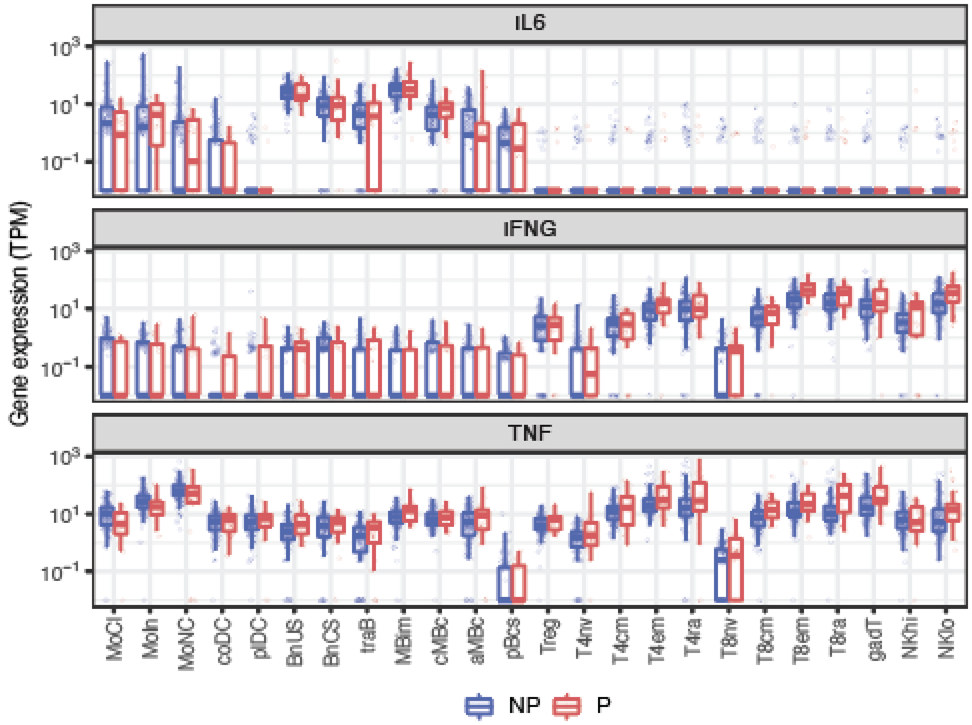


**Fig. S5. RNA gene expression levels of *IL6*, *IFNG*, and *TNF* in Non-progressors and Progressors.** *IL6*, *IFNG*, and *TNF* gene expression levels are displayed as transcripts per million (TPM) by cell subset for Non-progressors (NP) versus Progressors (P). Abbreviations for cell subsets are defined in table S3.

**Fig. S6.**

**A**

Wikipathways

| Pathway | fdr |
|---|---|
| *COMPLEMENT AND COAGULATION CASCADES* | 0.0046 |
| *MICROGLIA PATHOGEN PHAGOCYTOSIS PATHWAY* | 0.028 |
| *COMPLEMENT ACTIVATION* | 0.050 |

Reactome

| Pathway | fdr |
|---|---|
| *COMPLEMENT CASCADE* | 0.0063 |
| *CREATION OF C4 AND C2 ACTIVATORS* | 0.016 |
| *INITIAL TRIGGERING OF COMPLEMENT* | 0.050 |

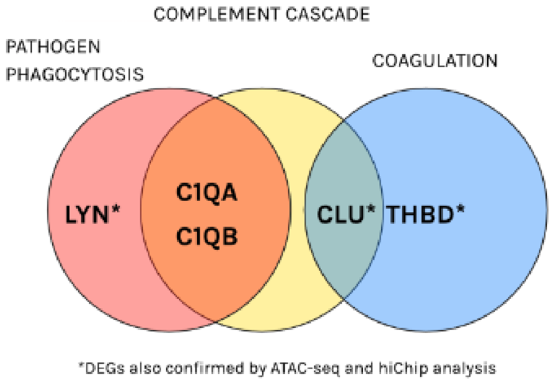


**B**

| Gene | Other studies of COVID-19 |
|---|---|
| *IL1R2* | Bost, 2022*(1)*; Reyes, 2020 *(2)* |
| *SAP30* | Ahren, 2021*(3)* |
| *HMGB2* | (on its homologue *HMGB1*; Chen R, 2020 *(4)*; Sivakorn, 2021 *(5)*; Passos, 2022 *(6)*; Ruskowski *(7)*) |
| *CLU**‡ | Singh, 2021 *(8)*; (in MERS-CoV Yeung, 2016 *(9)*) |
| *RNASE1**‡ | McDonald, 2021 *(10)*; Yoshida, 2022 *(11)* |
| *VSIG4* | Zhang, 2022 *(12)*; Yoshida, 2022 *(11)* |
| *S100A12* | Schulte-Schrepping, 2020 *(13)*; Silvin, 2020 (in neutrophils) *(14)*; Lei, 2021 *(15)* |
| *CD163** | Wendisch, 2021 *(16)*; Fassan, 2022 *(17)*; Gomez-Rial, 2020 *(18)*; Rajamanickam, 2021 *(19)*; Zingaropoli, 2021 *(20)*; Marocco, 2022 *(21)* |
| *C1QB* | Stephenson, 2021 *(22)*; Zhang, 2022 *(12)* |
| *MCEMP1** | van der Wijst, 2021 (in neutrophils) *(23)*; Carapito, 2021 *(24)* |
| *THBD**‡ | Gavriilaki, 2021 *(25)*; Asteris, 2022 *(26)* |
| *RAB13**‡ | (listed as capable of SARS-CoV-2 binding Tiwari, 2022 *(27)*) |
| *ASGR2* | Arunchalam, 2020 (appears in a list without further follow up) *(28)* |
| *HLX* | (appears in lists without further follow up Arunchalam, 2020 *(28)*; Knight, 2021 *(3)*) |
| *C1QA* | Ma, 2021 *(29)*; Stephenson, 2021 *(22)*; Zhang, 2022 *(12)* |
| *TRPM2* | Jaffal, 2021 *(30)* |
| *S100A9* | Schulte-Schrepping, 2020 (in neutrophils) *(13)*; Silvin, 2020 *(14)*; Mellet, 2022 *(31)* |
| *GPX1* | Saik, 2022 *(32)* (Seale, 2020 *(33)* found a protective role, contrary to our data) |
| *LYN**‡ | Rebendenne, 2022 *(34)* (in lung and colorectal epithelial cell lines) |
| *SVIL* | (Lyons-Weiler, 2020 *(35)*; Zhou, 2022 *(36)* as having direct interaction with SARS-CoV-2) |
| *MT-CO1* | Duan, 2022 *(37)*; McDonald, 2021 *(10)* |
| *LAIR2* | |

*DAR in ATAC-seq analysis ‡DAR in HiChip analysis

**Fig. S6. Pathway analysis and other COVID-19 literature on monocyte DEGs.** (**A**) Pathway analysis of combined monocyte differentially expressed genes (DEGs). Analysis was done for the Wikipathways and Reactome pathway sets (curated by the MsigDB collections) using a hypergeometric test of all monocyte DEGS with a FDR ≤ 0.1 cutoff. Venn diagram shows significant DEGs at a FDR ≤ 0.1 within the pathway sets. (**B**) Table of significant monocyte DEGs at a FDR ≤ 0.1 with references to associated COVID-19 literature for each gene.

Fig. S7.

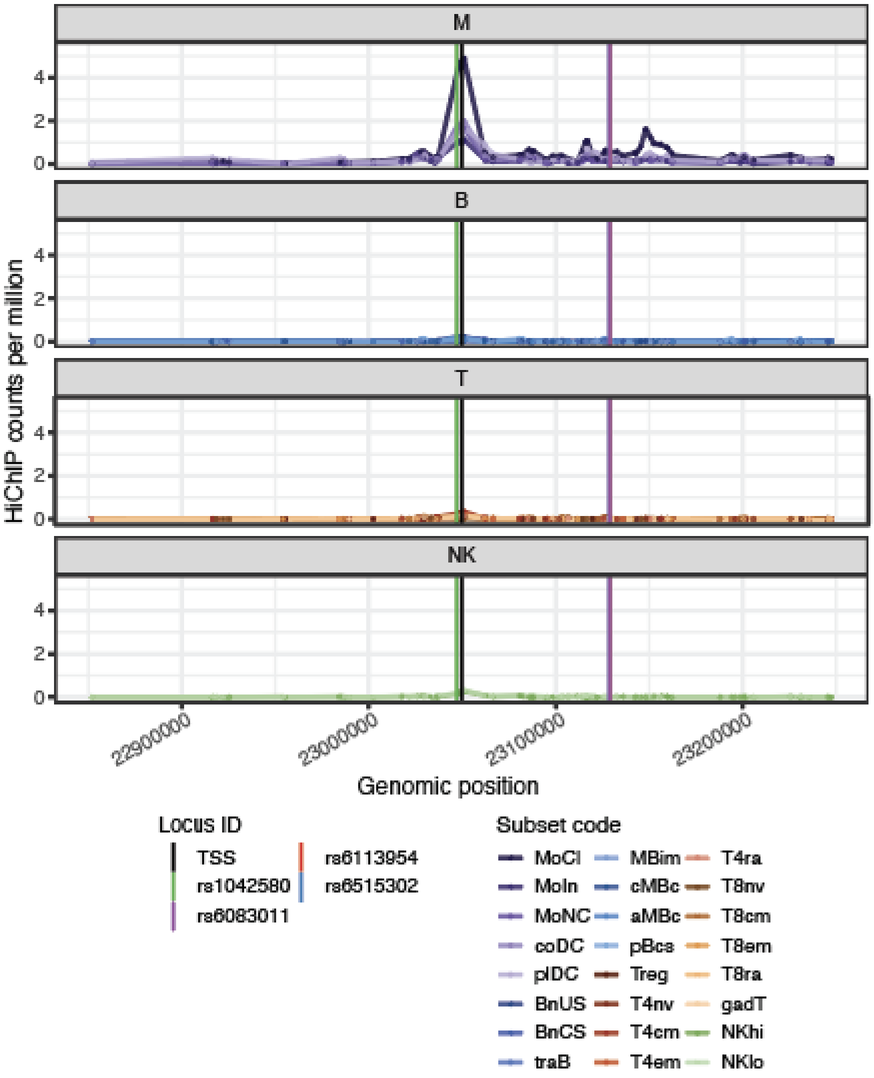


**Fig. S7. Chromatin contact via HiChIP around *THBD* in all subsets.** Chromatin contact via HiChIP between regions with eQTL SNPs and the THBD TSS in all cell subsets, grouped by B, M (monocyte), NK, and T cells. The black vertical line represents the transcriptional start site (TSS), and colored vertical lines represent variant positions, with the lines for rs6083011, rs6113954, and rs6515302 overlapping in the figure. Abbreviations for cell subsets are defined in table S3.

**Fig. S8.**

A

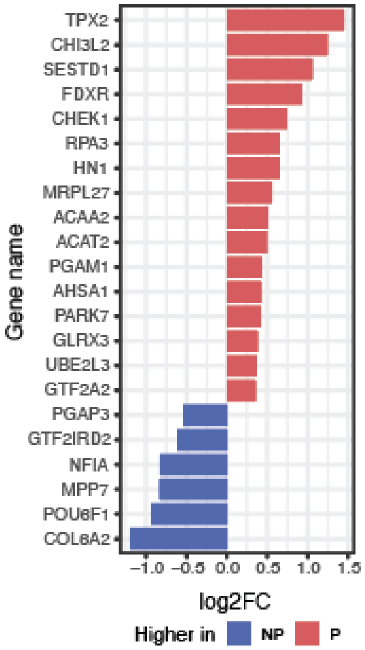


B

| Human Gene (with corresponding mouse gene in LCMV dataset) | Mouse Activation Model (LCMV) (ImmGen RNA-seq) | Concordance between CPR & LCMV |
|---|---|---|
| *TPX2, FDXR, CHEK1, RPA3, MRPL27, ACAT2, PGAM1, PARK7, UBE2L3, HN1, GTF2A2* | ↑ in LCMV | Yes |
| *PGAP3, MPP7, GTF2IRD2* | ↓ in LCMV | Yes |
| *COL6A2, NFIA, POU6F1* | ↑ in LCMV | No |
| *SESTD1, GLRX3, ACAA2, AHSA1* | ↓ in LCMV | No |

**Fig. S8. Univariate differential gene expression analysis in effector memory CD8+ T cells.** (**A**) Log2 fold-change (log2FC) of DEGs in the effector memory CD8+ T cells between Non-progressors (NP) and Progressors (P) at FDR ≤ 0.05. (**B**) DEGs in effector memory CD8+ T cells detected at FDR ≤ 0.05 overlapping with mouse gene equivalents in an lymphocytic choriomeningitis virus (LCMV) model of T cell activation.

### SUPPLEMENTARY MATERIALS AND METHODS

#### TaPE-seq OligoAb cocktail

The OligoAb cocktail includes a panel of oligo-barcoded antibodies enabling protein abundance estimation using Targeted Protein Estimation by sequencing (TaPE-seq). The markers targeted by the antibodies in the cocktail are listed in the tables below.

| Lineage | | | |
|---|---|---|---|
| CD3 | CD19 | IL2RA (CD25) | NCAM1 (CD56) |
| CD4 | CR2 (CD21) | IL7R (CD127) | TRDC; TRGC1 (gdTCR) |
| CD8A | FCGR2A/B (CD32) | ITGAX (CD11c) | |
| CD14 | FCGR3A/B (CD16) | MME (CD10) | |

| Functional | | | |
|---|---|---|---|
| ADGRG1 (GPR56) | CD47 | ICAM1 (CD54) | LILRB2 (CD85d) |
| ANPEP (CD13) | CD5 | ICOS (CD278) | LY9 (CD229) |
| B3GAT1 (CD57) | CD6 | ICOSLG (CD275) | CD60a (GD3) |
| BTLA (CD272) | CD69 | IL21R (CD360) | NCR1 (CD335) |
| CCR10 | CD7 | IL2RB (CD122) | NCR2 (CD336) |
| CCR2 (CD192) | CD70 | IL4R (CD124) | NT5E (CD73) |
| CCR4 (CD194) | CD80 (B7-1) | IL6R (CD126) | PDCD1 (PD-1) |
| CCR5 (CD195) | CD81 | ITGA1 (CD49A) | PDCD1LG2 (PD-L2) |
| CCR6 (CD196) | CD84 (SLAMF5) | ITGA2 (CD49B) | PECAM1 (CD31) |
| CCR7 (CD197) | CD86 (B7-2) | ITGA4 (CD49D) | PTGDR2 (CD294) |
| CD160 | CD9 | ITGA5 (CD49e) | PTPRC (CD45RA) |
| CD1A | CTLA4 (CD152) | ITGAE (CD103) | PTPRC (CD45RO) |
| CD1C | CX3CR1 (CD183) | ITGAL (CD11a) | PVR (CD155) |
| CD2 | CXCR2 (CD182) | ITGAM (CD11b) | SELL (CD62L) |
| CD200 | CXCR3 (CD183) | ITGB1 (CD29) | SIGLEC7 (CD328) |
| CD226 | CXCR4 (CD184) | ITGB2 (CD18) | SIGLEC9 (CD329) |
| CD244 (2B4) | CXCR5 (CD185) | KLRB1 (CD161) | SLAMF7 (CD319) |
| CD27 | DPP4 (CD26) | KLRD1 (CD94) | SLC3A2; SLC7A5 (CD98) |
| CD274 (PD-L1) | ENTPD1 (CD39) | KLRG1 | TIGIT |
| CD276 (B7-H3) | FASLG (CD178) | KLRK1 (NKG2D) | TNFRSF14 (CD270) |
| CD28 | FCRL3 (CD307c) | L1CAM (CD171) | TNFRSF18 (GITR) |
| CD38 | FCRL6 | LAG3 (CD223) | TNFRSF4 (OX40) |
| CD40 | HAVCR2 (TIM-3) | LAIR1 (CD305) | TNFRSF8 (CD30) |
| CD40LG (CD154) | HLA-ABC | LGALS3 (Galectin3) | TNFRSF9 (41BB) |
| CD44 | HLA-DRA | LILRB1 (CD85j) | VTCN1 (B7-H4) |

#### Multi-omic machine learning modeling

Block sparse partial least squares discriminant analysis (block.sPLS-DA) was used in identifying correlated feature scores between RNA-sequencing (RNA-seq) and assay for transposase-accessible chromatin using sequencing (ATAC-seq) data that are associated with progression status. Nested cross-validation (CV) was used, where the outer CV is a leave-one-out (LOOCV) and the inner CV performs a 5-fold grid search to identify the best hyperparameters using the validation data's area under the ROC curve (AUC). The hyperparameters used in the grid search were keepX and the design matrix covariance weighting between RNA and ATAC blocks. The top subsets were selected using the outer LOOCV AUC, and top features were taken from a model retrained on all data using the optimal hyperparameters.

#### Expression quantitative trait loci

To link genes to genome-wide association studies (GWAS) risk variants, we identified significant expression quantitative trait loci (eQTLs).

cis-eQTLs were generated using Matrix eQTL using multiple covariates, including sex, probabilistic estimation of expression residuals (PEER) factors derived from gene expression, and genotype principal components. Analysis was limited to variants within 250 kb of genes. eGenes were considered significant at a false discovery rate (FDR) ≤ 0.1, with this FDR being corrected after filtering eQTLs for genes that were differentially expressed genes (DEGs) for progression outcome at significance FDR ≤ 0.1.

#### Genetic associations

The association between genetic variants and progression was tested by regressing progression outcome on genotype, adjusting for age, sex, race, an indicator for comorbid respiratory conditions, and 10 principal components derived from whole genome sequencing data.

#### Literature review

Literature review of COVID-19 associated studies of DEGs was conducted using the Google search engine and defining the search terms as "gene name" + covid19 OR sars-cov-2. Where no literature was uncovered, an additional query of '"gene name" + immunology OR respiratory disease' was performed. Where results were published in similar cell types to those associated with the DEGs identified in our study, we reported those publications (**fig. S6B**). In cases where findings were only available in different cell types, we included annotations for these publications along with the cell type observed (**fig. S6B**).
